# StaBLE: digital metrics capture balance performance across a wide spectrum of balance tasks and abilities

**DOI:** 10.64898/2026.09.16.750744

**Authors:** Hannah Heigold, Parker S. Ruth, Julie Muccini, Dawit Lee, Sydney Barta, Ariana Rodrigues, Trevor Hastie, Kristen K. Steenerson, Scott L. Delp

**Affiliations:** Department of Bioengineering, Stanford University, Stanford, CA, USA; Department of Computer Science, Stanford University, Stanford, CA, USA; Department of Radiology, Stanford University, Stanford, CA, USA; Department of Statistics, Stanford University, Stanford, CA, USA; Department of Biomedical Data Science, Stanford University, Stanford, CA, USA; Department of Otolaryngology (Head and Neck Surgery), Stanford University, Stanford, CA, USA; Department of Neurology and Neurological Sciences, Stanford University, Stanford, CA, USA; Department of Mechanical Engineering, Stanford University, Stanford, CA, USA; Department of Orthopaedic Surgery, Stanford University, Stanford, CA, USA

## Abstract

Balance is fundamental to mobility and independence, yet it remains difficult to measure in a way that is both clinically meaningful and feasible for widespread adoption. There is a need for more quantitative, objective, and accessible balance assessments. To this end, we created the Stanford Balance Level Evaluation (StaBLE), a battery of 18 balance-challenging tasks and accompanying digital performance metrics computed from video that aimed to capture widely ranging balance abilities across 180 participants. We validated the StaBLE score against existing clinical scales and found that it correlated with age (ρ = -0.76, p<0.001), the Activities-Specific Balance Confidence Scale (ρ = 0.60, p<0.001), a Mini-Balance Evaluation Systems Test task (τ=0.55, p<0.001), 4-Stage Balance Test (τ = 0.55, p<0.001), and Short Physical Performance Battery (τ =0.57, p<0.001). StaBLE overcame ceiling effects in these existing scales and had a different underlying distribution for those identified at risk of falling and not (KS: 0.67, p<0.001). To reduce the time it takes to perform the test, we identified subsets of balance tasks that best predicted the StaBLE score across different populations. Together, our dataset, balance assessment protocol, and performance-based score demonstrate the validity of digital measures of balance.

## INTRODUCTION

Maintaining balance is essential for human movement, yet its importance is often only recognized when balance fails. A loss of stability without adequate recovery results in a fall, which can have devastating consequences, particularly for older adults. Unintentional falls are the leading cause of injury-related death in older adults and have substantial societal and economic costs.^1,2^ While falls are a prominent outcome of poor balance, balance itself is a multi-dimensional spectrum that influences how people of all ages and abilities move through the world and how athletes achieve peak performance. Fortunately, balance can be improved through targeted interventions, including strength training, sensory integration, and vestibular rehabilitation.^3–5^ Determining the extent to which these interventions improve balance relies on accurate assessment of balance ability.^6^

Balance remains difficult to quantify with methods feasible for widespread adoption. As a result, clinical screening often relies on questions about retrospective fall history. ^7^ These screenings are easy to deploy but are unsuitable to prevent a first fall or provide mechanistic insight. Clinical assessments of balance, deployed in physical therapy or specialist settings, use targeted tasks such as single-leg stance or eyes-closed standing.^8^ The Mini-Balance Evaluation Systems Test (Mini-BESTest),^9,10^ the CDC’s STEADI 4-Stage Balance Test,^11,12^ and the Short Physical Performance Battery (SPPB)^13,14^ are widely used, validated for fall risk assessment, and feasible to administer in minutes. However, these assessments rely on coarse scoring systems and exhibit ceiling effects, limiting their sensitivity. In addition, they depend on subjective evaluation by trained clinicians, which can limit standardization and scalability. Laboratory-based measures of kinematics successfully capture balance with precision,^15–17^ but remain impractical for clinical translation due to cost and expertise requirements.

Advances in digital health tools for measuring human movement present an opportunity to develop a quantitative and objective balance assessment that overcomes these limitations. Smartphone-based motion capture technologies expand access to whole-body human movement measurements with comparable quality to laboratory settings.^18–20^ The ubiquity of smartphones makes it plausible for widespread use. Prior work has demonstrated that models trained on video-derived pose data can predict clinical balance scores,^21–24^ highlighting the potential to extract meaningful information from complex movement patterns. However, these approaches typically replicate existing clinical scales and therefore inherit their limitations in sensitivity. The remaining challenge is how to leverage the unique capabilities of smartphones as powerful digital health tools to produce new measures of balance that complement existing clinical metrics, while maintaining interpretability.

To this end, we created the Stanford Balance Level Evaluation (StaBLE), a battery of balance-challenging tasks and accompanying digital performance score designed to capture a wide range of balance abilities. First, we established a repeatable protocol to collect video data and characterized the StaBLE score in participants with widely ranging balance abilities. Second, we validated the StaBLE score against existing clinical scales and tested that it removes ceiling effects. Third, to reduce the time of the assessment, we explored which subsets of balance tasks are most informative of balance ability across different populations.

## RESULTS

### Participants

We recruited 180 participants (117 female, 63 male) with diverse ages (18-87), diagnoses (71 self-reported), and competitive sports background (30 sports) to complete the StaBLE protocol. This resulted in a wide spectrum of balance abilities, ranging from individuals with impaired balance (i.e., vestibulopathy, peripheral neuropathy, Parkinson’s disease) to elite balancers (professional ballerinas, acrobats, or collegiate gymnasts). We collected data in 6 locations: the Stanford Human Performance Lab, the Stanford Ear Institute, three dance studio and theatre settings, and an adaptive physical education community center.

### StaBLE protocol

The StaBLE protocol comprised 18 tasks (**Sup. Table 1**) derived from common clinical assessments including standing tasks, dynamic tasks (i.e., jumping and reaching), and gait tasks. Five tasks included an eyes-closed component. The tasks were recorded on OpenCap^18^ using a standardized data collection setup with three smartphone cameras and a laptop computer, standard floor markings, and background drop cloths (**Fig. 1**). Data collection was performed with 30 minutes to set-up the room and perform motion capture calibration, less than 10 minutes to prepare each participant (donning socks, providing instructions, and collecting a neutral pose), and 25-35 minutes to complete the movement trials. Data collection could be performed by a single experimenter.

**Fig. 1.**
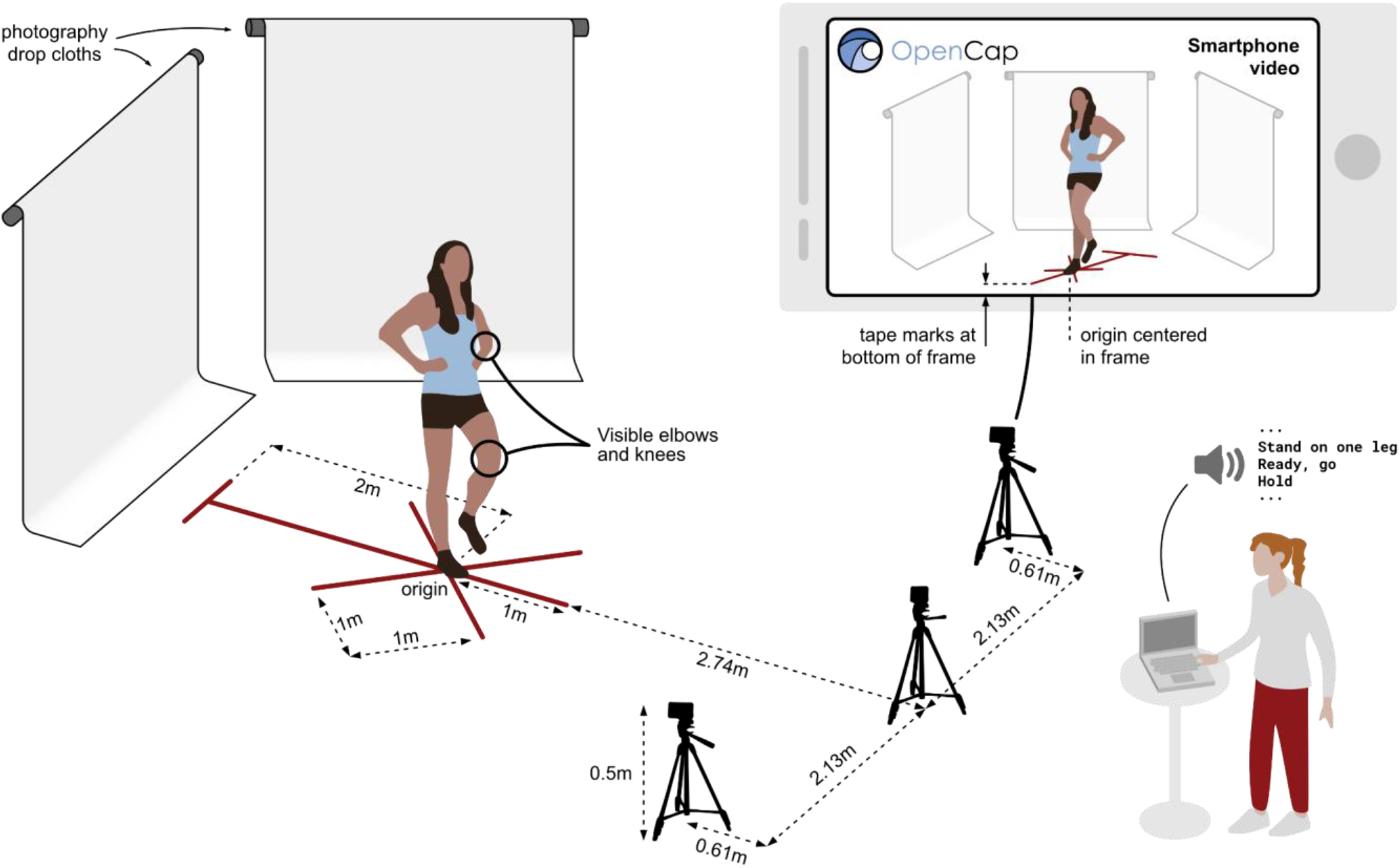
StaBLE data collection. Tape marks on the floor indicated the position and orientation for each task. Three tripods were positioned at a defined distance from the tape at a height of 0.5 m, each with a smartphone in landscape orientation. The camera field of view was standardized relative to the tape marks, horizontally centered on the origin, and vertically angled with the tape at the bottom of the view. All devices were connected to power and Wi-Fi (not shown). Participants were asked to wear clothing showing their knees and elbows, if possible. They were supplied with non-slip socks. Drop cloths were used to minimize background visual noise. We ran the OpenCap web app from a laptop computer, which also provided recorded audio cues for the standing and functional reach tasks.

### StaBLE score

We collected 3322 motion trials as 180 participants performed up to 18 different tasks. For safety, 98 trials across 24 participants were not performed (e.g., the Y-Balance and Jump tasks were not performed if an individual could not safely stabilize on one leg). We constructed 76 digital features to measure the participants’ degree of success in achieving the task objectives (**Sup. Table 2**). For example, in the Rise-to-Toes task, participants were instructed to rise to their toes as quickly as possible, as high as possible, and to hold for 5 seconds. These objectives were measured with the peak vertical center of mass speed, peak plantar flexion, and time the center of mass was above 50% of its peak vertical position (**Fig. 2a**). The features were aggregated by task into 18 task scores, which were further aggregated into a single StaBLE score, with higher scores representing better performance (**Fig. 2a**). We show this in exemplar videos across the range of StaBLE scores (**Fig. 3**). The StaBLE score had a left-tailed distribution (min: 0.08, max: 0.84, mean: 0.64, standard dev: 0.12) (**Fig. 2c**). Task scores display unique distributions with more challenging tasks tending to have lower means and higher standard deviations (**Fig. 2b**). This is seen in the progressive standing tasks: Standing Eyes-Closed (mean: 0.90, std: 0.08), Semi-Tandem (mean: 0.85, std: 0.10), Tandem (mean: 0.80, std: 0.17), Single-Leg-Stance (SLS) Eyes-Closed (EC) (mean: 0.81, std: 0.19), SLS Rise-to-Toes (RTT) (mean: 0.57, std: 0.21), SLS RTT EC (mean: 0.58, std: 0.22). The scores do not show ceiling effects. For tasks that could not be performed by everyone (SLS-RTT-EC, Y-Balance, Jump), there are clusters at zero.

**Fig. 2.**
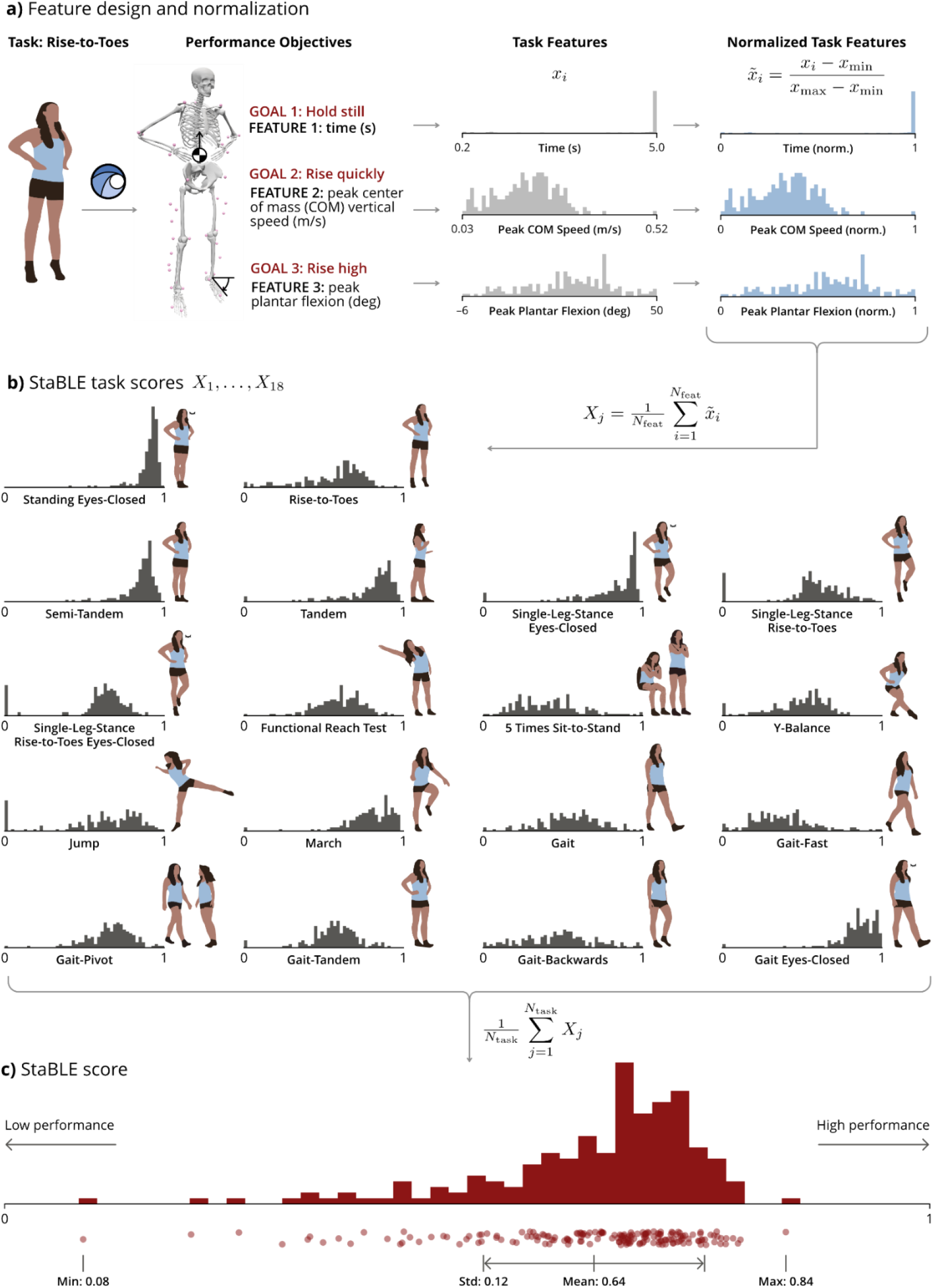
StaBLE tasks and score. **(a)** Task features convert task performance objectives into biomechanical features, normalized by the maximum and minimum value in the dataset. **(b)** Task scores aggregate normalized features. Distributions of task scores across 180 participants of diverse demographics and balance-related covariates are shown in the order performed left to right, top to bottom. Task score histograms share the same y-axis frequency scale. **(c)** Overall StaBLE score aggregates task scores from each of 18 balance-challenging tasks.

**Fig. 3.**
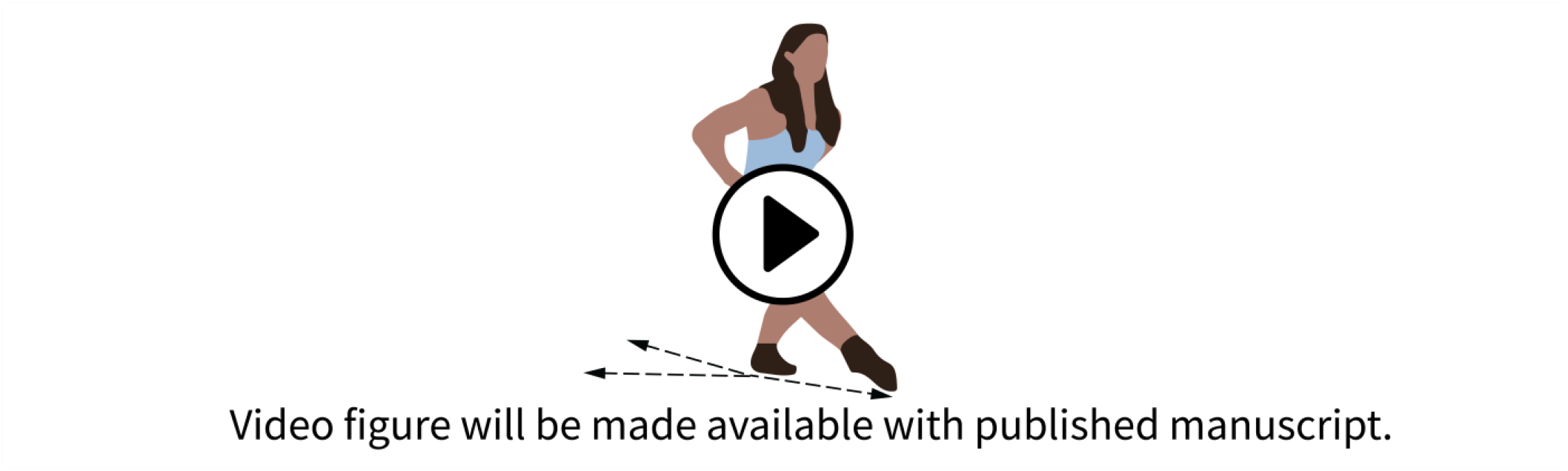
Video of representative tasks. Participants with StaBLE scores of 0.20 (0:00), 0.30 (0:24), 0.43 (0:58), 0.50 (1:37), 0.61 (2:24), 0.70 (3:05), and 0.84 (3:49) performing a sample of StaBLE tasks. All videos are taken from the smartphone to the participant’s left. In the PDF version of this article, click anywhere on the figure or caption to play the video in a separate window.

### StaBLE score validity

The StaBLE score had a strong negative correlation with age (ρ: -0.76, p<0.001) (**Fig. 4a**) and a positive correlation with balance self-efficacy measured by the Activities-specific Balance Confidence (ABC) Scale ^25^ (ρ: 0.6, p<0.001) (**Fig. 4b**). The ABC Scale is commonly used as an outcome measure and fall risk assessment tool ^26^, but is primarily validated in older and patient populations. In our dataset the ABC Scale had a ceiling effect with 39 maximal scores, whereas zero participants scored a maximum score of 1 on the StaBLE score.

**Fig. 4.**
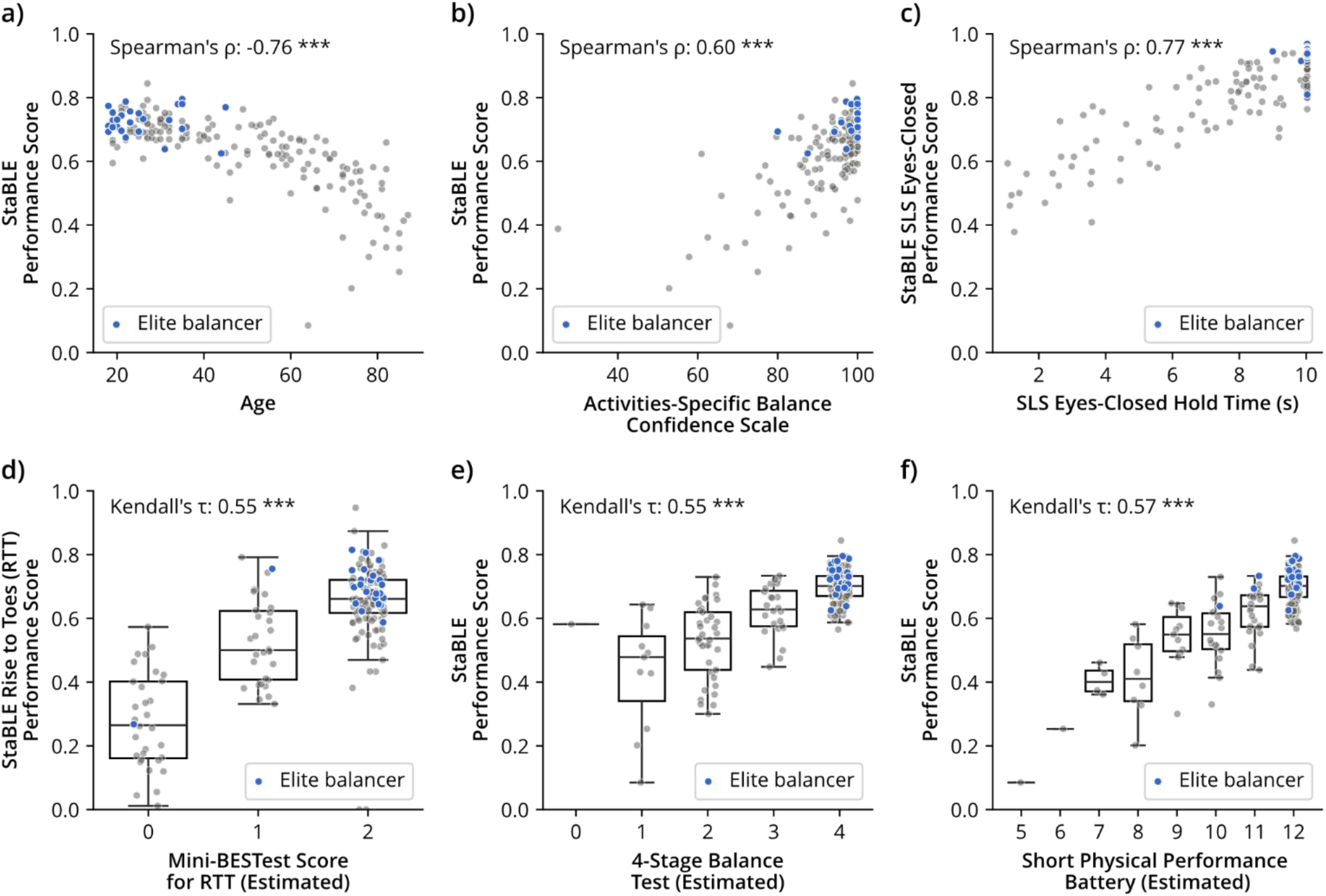
StaBLE and balance-related validation measures. The StaBLE score correlates with age and existing balance-related functional scales. Elite balancers are defined as participants who competitively train in gymnastics, dance, or acrobatics. **(a)** The StaBLE score decreases with age. **(b)** The StaBLE score is positively correlated with the Activities-specific Balance Confidence (ABC) Scale. Many participants reported maximal scores for the ABC Scale of 100 (n=39), but no participants achieved a perfect StaBLE score of 1. **(c)** The StaBLE Single-Leg-Stance (SLS) eyes-closed score is correlated with the eyes-closed time measures. By including sway metrics, the StaBLE task score overcomes the ceiling effect seen in time measures of single-leg stance with eyes closed. **(d)** The StaBLE Rise-to-Toes (RTT) score is positively correlated with the estimated Mini-BESTest task score. StaBLE scores are continuous rather than categorical, demonstrating greater spread in elite balance performance (std = 0.04) compared to Mini-BESTest which scored 24/26 elite balancers a maximal value of 2. **(e)** The total StaBLE score correlates with 4-Stage Balance Test and **(f)** Short Physical Performance Battery scores estimated from smartphone video. (*** p<0.001)

Among the simplest and most common tests of balance is hold time, e.g. the ability to stand on one leg with eyes closed for ten seconds. In our dataset, ten participants reached the test ceiling by holding the stance for a total of ten seconds across the right and left leg. Since the StaBLE score for this task combined hold time with metrics of sway and stability, it correlated closely with pure hold time (ρ = 0.77, p<0.001) and eliminated ceiling effects (no participants achieved perfect task scores of 1) (**Fig. 4c**).

The Mini-BESTest,^9,10^ 4-Stage Balance Test,^11,12^ and SPPB ^13,14^ included tasks that are also part of the StaBLE balance protocol and are evaluated with ordinal scales. One example is the Rise-to-Toes task, which the Mini-BESTest scores on an ordinal 3-point scale based on success in holding the task, observable stability, and maintaining plantarflexion. The continuous StaBLE Rise-to-Toes score was correlated with the estimated Mini-BESTest Score for this task (τ = 0.55, p<0.001) (**Fig. 4d**). Out of the 26 elite balancers, 24 scored the maximum value on the Mini-BESTest assessment, indicating a strong ceiling effect. These participants had a range of non-maximal StaBLE Rise-to-Toes scores (std: 0.04), demonstrating StaBLE may better differentiate balance at the higher end of performance. The total StaBLE score was correlated with the estimated 4-Stage Balance Test (τ = 0.55, p<0.001) (**Fig. 4e**) and the estimated Short Physical Performance Battery (SPPB) (τ = 0.57, p<0.001) (**Fig. 4f**) scores. The StaBLE score included additional tasks beyond the progressive standing, gait, and sit-to-stand tasks in these ordinal scales, accounting for some discrepancy. StaBLE overcame the ceiling effects in the 4-Stage Balance Test (107 participants scored maximally) and SPPB (113 participants scored maximally) by extending the difficulty of the tasks and including the sway metrics for standing tasks. The StaBLE score also revealed substantial difference in balance ability among participants with the same score for the Mini-BESTest Rise to Toes Score, the 4-Stage Balance Test, and SPPB, suggesting greater capacity to differentiate balance ability.

Falls are an impactful outcome of poor balance. Therefore, a balance metric should have a different underlying distribution for those at risk and not at risk of falling. We found a significant difference in the distribution of StaBLE scores between participants identified as at risk of falling and not, as determined by the CDC STEADI Fall Risk criteria^11^ (KS: 0.67, p<0.001) (**Fig. 5a)**. Every individual task score in StaBLE, except Gait Eyes-Closed, had a significant difference in distribution between fall-risk and not-at-risk groups (**Sup. Fig. 1**). The task scores with the greatest difference in distribution were Single-Leg-Stance Rise-to-Toes Eyes-Closed (KS=0.65, p<0.001), Single-Leg-Stance Rise-to-Toes (KS = 0.62, p<0.001), Y-Balance (KS = 0.62, p<0.001), and Gait-Pivot (KS = 0.61, p<0.001). There was also a significant difference in distribution of StaBLE scores between participants who had a history of falls compared to those who did not (KS: 0.44, p <0.05) (**Fig. 5d**). While the estimated 4-Stage Balance Test scores and SPPB scores had a significant difference in distribution between those at risk for falling and not (**Fig. 5b, Fig. 5c**), there was no significant difference in distribution between those who had a history of falls and those who did not (**Fig. 5e, Fig. 5f**).

**Fig. 5.**
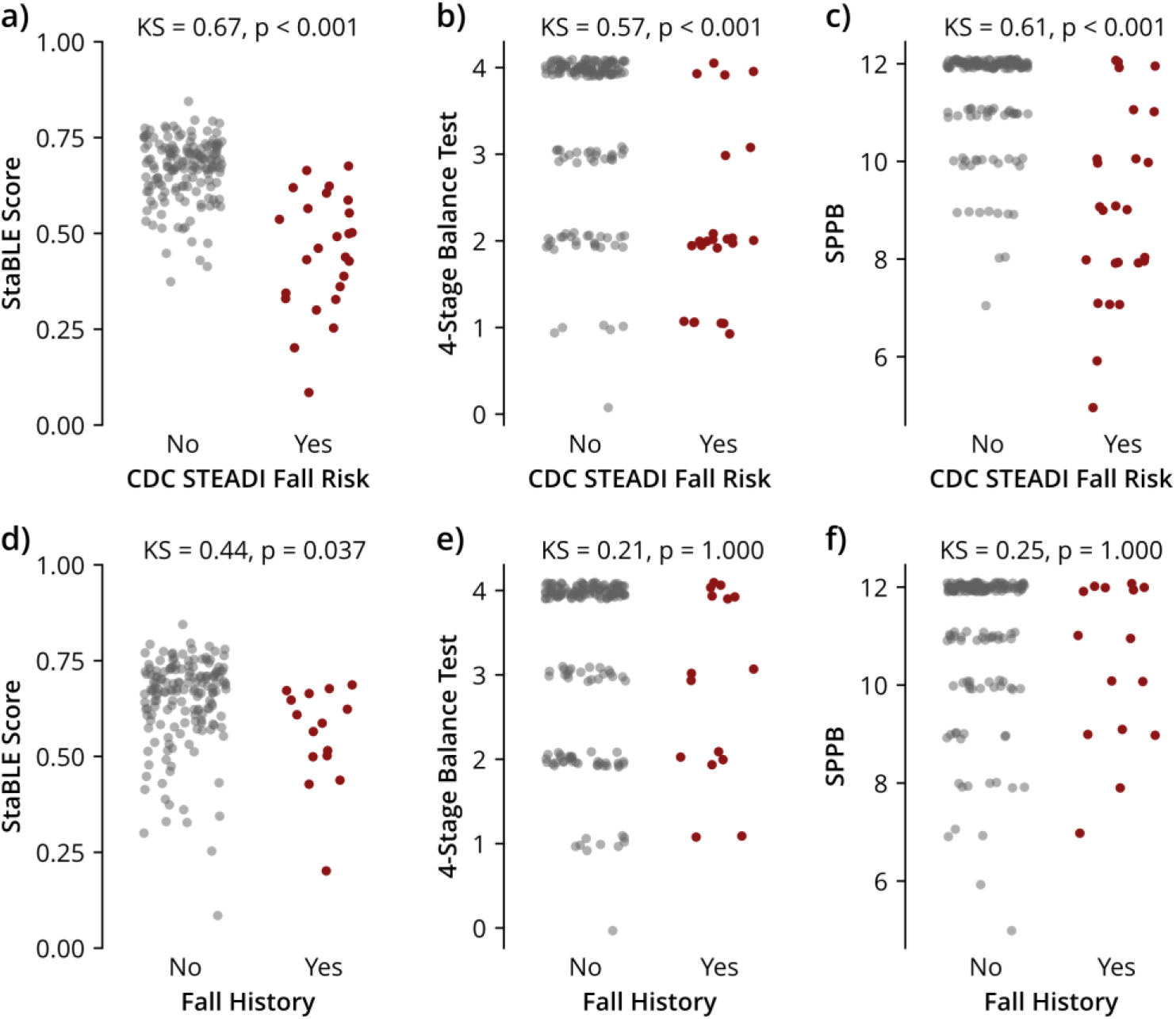
StaBLE and fall-related validation measures. **(a)** Participants identified as fall risk (CDC Stay-Independent 12-question tool score of 4 or greater, n = 25) had a different underlying distribution of StaBLE scores compared to those not identified at risk. **(b)** This statistical difference in distribution was also seen in the estimated 4-Stage Balance Test and **(c)** Short Physical Performance Battery (SPPB) scores. **(d)** Only the StaBLE score had a statistically different distribution for those with a history of falls (n = 15) and those who did not. **(e)** 4-Stage Balance and **(f)** SPPB did not have a statistical difference for fall history. All comparisons were made with the Kolmogorov-Smirnov (KS) test, applying a Bonferroni correction.

### Task subset selection

To explore whether it was possible to reduce the time it takes to assess a participant, we used forward subset selection^27^ to identify a subset of tasks that could reproduce the StaBLE score in a linear model (**Fig. 6a**). Y-Balance was the first task to be selected in the model. This task alone explained 63% of the variance in the StaBLE score, estimated with cross-validated coefficients of determination (R^2^). The model subsequently selected Single-Leg-Stance Eyes-Closed, Gait-Backwards, Gait-Pivot, and Single-Leg-Stance Rise-to-Toes Eyes-Closed. These five tasks could explain 88% of the StaBLE score variance. Despite the heterogeneity of balance tasks, these results indicated there is shared information across the performance of the tasks, allowing for a reduced-task assessment.

**Fig. 6.**
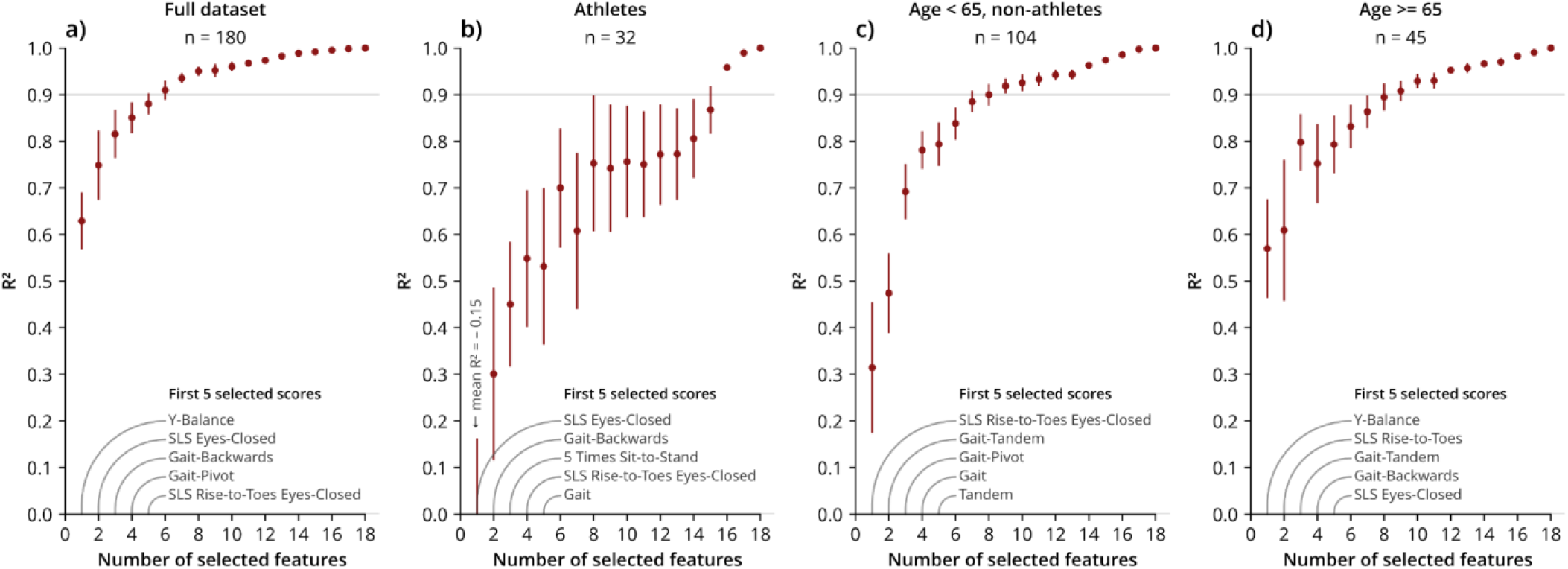
StaBLE task importance. Forward subset selection of task scores predicting the StaBLE score across the **(a)** full dataset **(b)** current competitive athletes, **(c)** adults under 65 years old who are not athletes, and **(d)** adults 65 and older. The mean R2 across the folds for each model size is plotted as a point with a bar for the standard error of the R2 across the folds for each model size. The first 5 selected scores were those chosen after five task scores were added to the model using the full dataset. (SLS = Single-Leg-Stance)

Unique subsets of tasks were selected when the analysis was performed on different groups. Athletes had the lowest variance in StaBLE scores (mean: 0.71, std: 0.07). Within this low-variance group, there was a lower correlation between task scores, resulting in a greater number of tasks required to explain 90% of the variance. **(Fig. 6b)**. A strength-dominant task, 5 Times Sit-to-Stand, was included among the first selected tasks. For adults under 65 (non-athletes) (**Fig. 6c**) and adults over 65 (**Fig. 6d**), 8 tasks explained 90% of the variance. All groups included gait and standing balance tasks in the first five selected tasks with Single-Leg-Stance Eyes-Closed, Gait-Backwards and Single-Leg-Stance Rise-to-Toes Eyes-Closed being selected the most frequently. Any combination of these five tasks could be performed in under 10 minutes.

## DISCUSSION

The StaBLE score, computed from video of balance-challenging tasks across a cohort with diverse balance abilities, addressed limitations of existing scales and demonstrated the potential of digital balance metrics. StaBLE does not require specialized operators or equipment to compute scores of balance performance, making it an accessible and objective assessment. Our dataset enabled data-driven selection of informative tasks and identified tasks that differ from common practice. The StaBLE score extended the capabilities of existing clinical scales by differentiating high-performance balance.

We developed a method for high-quality video-based data collection that could be completed with a single person. Markerless motion capture with OpenCap enables scalable data collection.^18,28^ This allowed us to collect data (n=180) in six unique locations, including dance studios and community centers, leading to a dataset that includes participants who would not typically visit a laboratory. We demonstrated that OpenCap can perform robustly across data collection settings. Intentional design of camera orientation, environment set-up, and participant clothing, along with recorded cues and standardized instructions, enabled consistent data collection. We include our optimized set-up for standardization and reproducibility.^29^ Since the StaBLE score is computed from kinematic data, it can be generalized to alternate motion capture tools that produce kinematics, thus avoiding reliance on a specific data capture modality. The size of the dataset was sufficient to apply data-driven methods, establish normative trends (**Fig. 4a**), and design and validate generalizable balance metrics.

We recruited participants with varied balance abilities, ranging from impaired to elite balance performance. Many existing digital measures of balance have been developed to separate healthy and impaired groups, either recruiting from a specific disease or fall-risk population^30,31^ or focusing on older adults.^24,29^ Byincluding high-performance balance abilities, StaBLE better approximated the balance performance ceiling and overcame the artificial ceiling effect imposed by many clinical scales.^9,14,25^ This could lead to improved sensitivity to changes in higher balance performance.

We designed the StaBLE protocol to include a large battery of balance-challenging tasks, rather than a single task or clinical scale. We selected tasks to challenge different facets of balance, including strength, limits of stability, anticipatory postural adjustments, gait, cognitive processing, and sensory reweighting,^9,12,14,32^ and to challenge varied ability levels. This included tasks typically reserved for athletic assessment (i.e., Y-Balance/modified star excursion^33–36^ and single-leg jump landing^37,38^) and extended tasks, such as adding a rise-to-toes and eyes-closed component to the single-leg-stance (**Fig. 2**). With our battery of tasks, we demonstrated the robustness of our single set-up (**Fig. 1**). We reduced occlusion and potential data loss with camera location and orientation and task orientation (i.e. instructing participants to turn to a 45 degree angle during Tandem to improve motion tracking^24^). By starting with a larger set of tasks, we could apply a data-driven approach for task selection, potentially enabling new insight on the optimal balance assessment tasks and the relationships between tasks (**Fig. 6**). This approach selected Y-Balance as a task that best captured overall balance abilities, a task which is typically not included in fall risk assessments.^11,24,29,31^ Given the unique set of tasks chosen for each group (**Fig. 6**), we envision alternate subsets of tasks could be selected based on specific application areas and future studies.

The StaBLE score is interpretable and overcomes ceiling effects of existing scales. To circumvent the challenges of defining a direct measure of balance or anchoring the StaBLE score in existing balance-related measures,^24^ which are potentially biasing or limiting, we designed the StaBLE score to measure *performance* on a battery of balance-challenging tasks. The StaBLE score uses features that directly measure the degree of success in achieving the objectives of each task, which is something that can be understood and computed from kinematic data (**Sup. Table 2**). This enabled dimensionality reduction of kinematic trajectories to lower dimension features and scores. Prior studies have selected features based on separating a specific disease population from healthy controls^30^ or to predict fall risk.^31^ Falls are an important outcome of poor balance, but there should be caution in using fall history as a direct label for balance. Falls have multi-factorial risk,^7^ are susceptible to reporting bias,^39^ and this label alone lacks the context surrounding a fall. For example, in our dataset we found 38 individuals reported a fall in the previous year with a mean age of 49 years old. Upon further inspection, 8 of these individuals were high-performing athletes, with some reporting 4 or more falls, likely encountered in their sport. While these falls may have implications for athletic performance or injury, they do not imply poor balance ability.

Exemplar videos reveal the effectiveness of the task scores and overall StaBLE score at capturing balance performance across a range of abilities (**Fig. 3**). The StaBLE score also agreed with anticipated relationships such as declining with age and correlating with balance confidence (**Fig. 4a, b**). Future work will explore the discordance between self-efficacy and measured ability via the StaBLE score and how this relates to real-world outcomes, such as prospective falls.^40^ The StaBLE score agreed with existing clinical scales (i.e. single-leg-stance hold time, Mini-BESTest, 4-Stage Balance Test, SPPB), but removed the ceiling effects, evidenced by the number of people achieving maximum scores. Several design choices enabled the StaBLE score to exhibit no ceiling.

Each individual feature was normalized based on the highest and lowest values among the 180 participants in this dataset. Since our participants had varied balance abilities, this resulted in a high ceiling and a low floor for each feature. Aggregating features and tasks further mitigated ceiling and floor effects. Achieving a perfect StaBLE score of 1 would require achieving the maximum possible performance on every feature for every task. These methods, along with the inclusion of difficult tasks, resulted in StaBLE showing no ceiling and floor effects (**Fig. 2c**). By removing ceiling effects, StaBLE is positioned to not only detect early-stage loss in balance ability but also to capture balance improvements from training. We show there is a difference in the underlying distributions of the StaBLE score between individuals at risk of falling and not, and those who self-reported 2 or more falls in the previous year (non-athletes) and those who did not (**Fig. 5**). With future work validating the score with prospective falls across various contexts and demographics, we anticipate StaBLE will be a valuable tool in fall risk assessment, a key application area for balance assessments.

There are several limitations of this work, notably in validation and score design. Further work is needed to characterize and validate the StaBLE score before it is implemented in clinical practice. The score has not been assessed for test-retest reliability, which will be important to determine the minimum detectable change and understand its sensitivity to training and repetition, time of day, and other factors known to influence balance performance.^41,42^ In addition to characterizing the variability within a person, future studies are required to understand the longitudinal response of the StaBLE score to balance training, injury, disease, and aging.

The StaBLE score makes simplifications that may miss nuances of the balance system needed for diagnosis and personalized intervention. The StaBLE score provides a snapshot measure of balance performance at a single point in time. Therefore, it may not reflect routine performance which can be influenced by medication, episodic symptoms, training, sleep, and other factors.^41–44^ It will be important to connect the score to real world balance performance and outcomes. The linear combination of features into the performance scores avoided the reliance on statistically learned embeddings, but assumed the features are additive and equally weighted. Future work should explore alternate strategies for aggregating features that account for factors such as progressive task difficulty. The StaBLE score is a single-dimensional score that represents overall balance ability. It can be used to identify a balance deficit, triggering follow-up, but cannot yet point to the underlying cause of the balance deficit. Further metric development is needed to characterize the subsystems and dimensions of balance for more specific diagnoses,^32^ beyond a task level score. One area of balance control we did not include was reactive balance. These assessments require equipment (i.e., a treadmill and harness) or additional evaluators, which did not meet our design constraints. Although further development is needed to make the StaBLE assessment accessible to clinical settings, this work provides a valuable baseline that will enable future work.

## METHODS

### Participants

Prior to participation, participants provided written informed consent to a protocol approved by the Stanford University Institutional Review Board (IRB-67713). Participants were included if they were 18 years or older, had not fallen in the past month, had no surgeries in the past 6 months, and were able to independently stand without support and walk 3 meters. We recruited broadly from the local community including various fitness studios, community centers, and clinics to ensure a range of balance abilities.

### StaBLE protocol

Participants completed the following survey series on RedCAP:^45,46^ demographics, medical history and medication, sport participation, International Physical Activity Questionnaire (IPAQ) Short Last 7 Days Self-Administered Format,^47,48^ fall history, CDC Stay Independent: a 12-question tool,^11^ Activities-specific Balance Confidence (ABC) Scale.^25^ Participants received the surveys within 48 hours before data collection and completed them before the balance protocol.

We designed the balance protocol to challenge the various subsystems of balance, remove ceiling effects, and require no specialized equipment or additional personnel in the capture window. We drew tasks from existing scales and extended them to challenge high-performance balance. We recorded the tasks using smartphone videos and the open-source platform, OpenCap (default settings, 60 fps, HRNet). We piloted multiple smartphone configurations and chose the one that minimized occlusion across diverse tasks with a single set-up. To standardize data collection, we had the same experimenter read scripted instructions to direct the protocol across all participants and we used an audio recording with cues for the standing tasks to ensure consistent timing. The participants only repeated a task if they did not follow instructions. For the jump task only, participants were allowed to practice before recording. For reliable video synchronization, we asked the participants to raise their right arm overhead at the beginning of each trial. We asked participants to wear shorts and a t-shirt or tank-top in contrasting colors (i.e. show their knees and elbows), to improve pose estimation. To control for variability due to shoe type, we supplied participants with non-slip socks.

### Data processing

We manually reviewed all 3322 motion trials and their OpenCap outputs for quality. For 30 trials (0.9%), OpenCap gave a solution with significant error due to multi-video synchronization failure or errors from occlusion. These trials were still included in the analyses. For 25 participants, there was a misalignment of the global and pelvis coordinate frame, likely introduced by user error during the calibration process.^18^ We applied a rigid transformation to the pelvis across all these participants’ trials. OpenCap failed to process only 3 trials (<0.1%) either due to failed video synchronization or user error.

### StaBLE score design

To design an objective balance performance score, we converted the goals of each task into features automatically segmented and computed from augmented marker data, joint kinematics, and musculoskeletal model output from OpenCap across all participants (**Sup. Table 2**). Some task goals were given explicitly to the participant, such as asking them to rise to their toes as quickly as possible, whereas others were implicit in the task, for example staying in a straight line while walking with eyes closed. For the 3 failed trials, the features were imputed using a linear regression with age for that feature. We normalized each feature by the maximum and minimum value in the dataset so that the best performance for that feature was assigned a score of 1 and the worst performance was assigned a 0. Features with the goal of maximization (e.g., speed) scaled relative to the minimum in the dataset

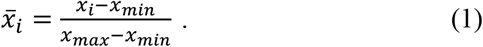

Features with the goal of minimization (e.g., deviation) scaled relative to the maximum in the dataset

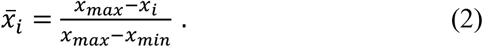

We assigned a normalized feature score of zero for participants unable to attempt or achieve a task. The score for each task is the mean of the normalized features for that task. For the jump task, if only one side was performed (e.g., due to injury), the jump score was the mean of the completed side features. The StaBLE score is the mean of the task scores.

### StaBLE score validity

We used Spearman’s rank correlation coefficient and Kendall rank correlation coefficient to compare the StaBLE score against continuous and ordinal metrics, respectively. The StaBLE Single-leg-stance (SLS) Eyes-Closed score, composed of sway and time features (succession of eyes open/closed right/left), was compared against the combined SLS eyes-closed hold time on both sides (**Fig. 4c**). We estimated clinical scale scores from the video data. Prior work has shown that video-based measures can reproduce timed function tests in clinical assessments.^28^ The Mini-BESTest Rise To Toes score was estimated using peak plantar flexion (range), standard deviation of the C7 virtual marker (sway), and hold time. For the 4-Stage Balance Test, we used hold time for the progressive standing tasks on the right side, applying stopping rules. Some thresholds were set to be below 10 seconds based on the cuing during the StaBLE assessment. These criteria were similarly applied to the SPPB and combined with times for the gait speed test and repeated chair stands.

We labeled participants as at risk for falling if they had a CDC Stay Independent: a 12-question tool score of 4 or greater.^11^ We compared the distribution of StaBLE, 4-Stage Balance Test, and SPPB scores for at-risk and not-at-risk participants using the Kolmogorov–Smirnov test. We also performed this test for individuals with and without a fall history. We defined fall history as participants who self-reported a history of 2 or more falls in the past year who were not currently participating in competitive sport or athletic performing arts. We applied a Bonferroni correction for multiple hypothesis testing to all statistics (**Fig. 5, Sup. Fig. 1**).

### Task subset selection

We used forward subset selection^27^ to iteratively add task scores to an ordinary least squares model predicting the total StaBLE scores. Tasks were selected with the greatest increase in explained variance (R^2^), using the full dataset. We obtained a cross-validated R^2^ with a K-fold cross validation with K=5 (**Fig.6**).

## DATA AVAILABILITY

The engineered features, clinical scores, participant demographics, and covariates for all participants are freely available at: https://doi.org/10.5281/zenodo.22286165

## CODE AVAILABILITY

The code used to compute the engineered features, perform statistical tests, and generate figures in this work is available in the following GitHub repository: https://github.com/stanfordnmbl/stable-analysis

## ACKNOWLEDGEMENTS

We gratefully acknowledge the contributions of our participants, data collection partners (Barbara McCarthy, M.S., Rachel Recinos Abair, M.S., Dr. Elizabeth Hutter, Psy.D., and Dr. Cherie Walker, PhD), the OpenCap development team (Scott Uhlrich, PhD, Antoine Falisse, PhD, Carmichael Ong, PhD, Matthew Petrucci, PhD, and Jennifer Hicks, PhD), and data collection assistants (Jimena Gutierrez, Kirsten Seagers, PhD, Kristen Steudel) for their contributions to this work.

This work was supported in part by the Joe and Clara Tsai Foundation through the Wu Tsai Human Performance Alliance at Stanford University and the U.S. National Institutes of Health (NIH) under Grants P41EB027060 and P50HD118632. We acknowledge the support of the Natural Sciences and Engineering Research Council of Canada (NSERC), [587678-2024]. Cette recherche a été financée par le Conseil de recherches en sciences naturelles et en génie du Canada (CRSNG), [587678-2024].

The funders played no role in study design, data collection, analysis and interpretation of data, or the writing of this manuscript

## AUTHOR CONTRIBUTIONS

HH, KKS, SLD conceived the project. SLD acquired funding. HH, JM, KKS, SLD designed the study protocol. JM managed the IRB. HH, JM, KKS recruited participants. HH led scheduling and data collection. JM, DL, SB, AR assisted with data collection. HH, SB, and AR performed data cleaning. HH designed features. HH and PSR designed the StaBLE score. HH wrote code for segmenting and extracting features. HH and DL performed feature quality assurance. TH advised on statistics and data analysis methods. HH and PSR performed data analysis and prepared figures. HH wrote the initial manuscript draft, PSR and SLD provided critical revisions to the manuscript, JM, DL, SB, AR, TH, KKS reviewed and edited the manuscript. All authors read and approved the final manuscript.

