## Supplementary Materials for "StaBLE: digital metrics capture balance performance across a wide spectrum of balance tasks and abilities"

### SUPPLEMENTAL MATERIALS

**Supplementary Table 1 | StaBLE task descriptions.**

| Task | Instruction | Timing<br>(Recorded Cues) | Sample of Clinical Measures with Task |
|---|---|---|---|
| **1. STANDING EYES-CLOSED**<br>*Standing-EC*<br>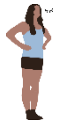<br>Stance with feet together, eyes open (10 seconds) and eyes closed (10 seconds). Hands on hips. | "We are going to do a sequence of standing tasks that will get progressively harder. Try to hold still in the task, but if you are unstable try to take a step rather than fall. When I say 'go', place your feet side-by-side, place your hands on your hips, look straight ahead. Then I will instruct you to close your eyes and hold that for 10 seconds." | Video record: 0s<br>"Arm Raise & Lower": 2s<br>"Go": 5s<br>"Eyes closed": 16s<br>"Neutral, Eyes Open": 26s<br>Stop recording: 28s | • Romberg Test<br>• Clinical Test of Sensory Interaction and Balance (CTSIB)<br>• 4-Stage Balance Test<br>• Mini Balance Evaluation Systems Test (Mini-BESTest)<br>• Short Physical Performance Battery (SPPB)<br>• Berg Balance Scale (BBS)<br>• Balance Error Scoring System (BESS) |
| **2. RISE-TO-TOES**<br>*RTT*<br>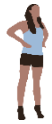<br>Rise to toes (5 second hold). Feet hip width. Hands on hips. | "Place your feet shoulder width apart. Keep your hands on your hips and look straight ahead. When I say 'go', try to rise as high as you can onto your toes, as quickly as possible, and hold this for 5 seconds until I say 'neutral'. " | Video record: 0s<br>"Arm Raise & Lower": 2s<br>"Hands on hips": 4s<br>"Go": 6s<br>"Hold": 7s<br>"Hold": 9s<br>"Neutral": 11s<br>Stop recording: 13s | • Mini-BESTest |
| **3. SEMI-TANDEM**<br>*Semi-Tandem*<br>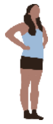<br>Semi-tandem stance, right and left leg (10 seconds). Hands on hips. | "Next, we will do semi-tandem stance. Have your feet shoulder width apart and your hands on your hips. Look straight ahead. When I say go, step your RIGHT foot forward so the LEFT big toe is in the middle of your foot, and your feet are glued together. After 10 seconds, I will ask you to stand neutral and then place the LEFT foot in front for 10 seconds. If needed you can look at your feet to place them, but then look straight ahead while holding" | Video record: 0s<br>"Arm Raise & Lower": 2s<br>"Hands on hips": 4s<br>"Go right": 6s<br>"Neutral": 17s<br>"Other side go": 19s<br>"Neutral": 30s<br>Stop recording: 33s | • 4-Stage Balance Test<br>• SPPB |

| Task | Instruction | Timing<br>(Recorded Cues) | Sample of Clinical Measures with Task |
|---|---|---|---|
| **4. TANDEM**<br>*Tandem*<br>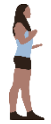<br>Tandem stance, right and left leg (10 seconds). Arms held with 90-degree elbow flexion, 45-degree external shoulder rotation. | "Now, when I say go, you will place your RIGHT foot all the way in front of your foot like you are on a balance beam with your big toe against your heel. For this task your arms will be in front, like you are holding a dinner plate. You will start on the right diagonal with your right foot in front. After 10 seconds, I will say, 'neutral face front' and then 'turn LEFT and step tandem', and we will repeat with the LEFT leg in front towards the LEFT diagonal. You may look down to check your position but then look up to hold" | Video record: 0s<br>"Arm Raise & Lower": 2s<br>"Arms in dinner platter": 4s<br>"Turn right": 6s<br>"Step tandem": 7s<br>"Face front": 18s<br>"Turn left": 20s<br>"Step tandem": 21s<br>"Neutral front": 32s<br>Stop recording: 35s | • 4-Stage Balance Test<br>• SPPB<br>• BBS<br>• BESS<br>• Sharpened Romberg |
| **5. SINGLE-LEG-STANCE EYES-CLOSED**<br>*SLS-EC*<br>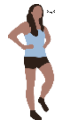<br>Single leg stance with eyes open (10 seconds) and eyes closed (5 seconds) on right and left leg. Hands on hips. | "Start in neutral. When I say 'go' you will lift your LEFT leg. Your leg can be whatever height you want but keep it off the ground and don't rest your leg on your standing leg. Keep your hands on your hips. After 10 seconds, I will ask you to close your eyes. Even if you put your leg down, try to keep your eyes closed and lift your foot back up for the next 5 seconds. I will let you know when to put your foot down and we will repeat lifting your RIGHT leg." | Video record: 0s<br>"Arm Raise & Lower": 2s<br>"Hands on hips": 4s<br>"Go": 6s<br>"Eyes closed": 17s<br>"Neutral, eyes open": 23s<br>"Other side go": 25s<br>"Eyes Closed": 36s<br>"Neutral eyes open": 42s<br>Stop recording: 45s | • 4-Stage Balance Test<br>• Mini-BESTest<br>• BBS<br>• BESS |
| **6. SINGLE-LEG-STANCE RISE-TO-TOES**<br>*SLS-RTT*<br>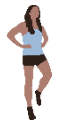<br>Single leg stance with rise-to-toe (5 seconds) on right and left leg. Hands on hips. | "Now, we are going to have you go onto your RIGHT leg again when I say 'go', but after three counts I will say 'rise' and you will rise to your toe. Your goal is to go as high as you can, without putting your LEFT foot down or your RIGHT heel down for 5 seconds. Then, I will instruct you to stand in neutral and we will repeat on the other side. Again, keep your hands on your hips." | Video record: 0s<br>"Arm Raise & Lower": 2s<br>"Hands on hips": 4s<br>"Go": 6s<br>"Rise": 8s<br>"Hold": 10s<br>"Neutral": 13s<br>"Other side go": 15<br>"Rise": 17s<br>"Hold": 19s<br>"Neutral": 22s<br>Stop recording: 25s | |

| Task | Instruction | Timing<br>(Recorded Cues) | Sample of Clinical Measures with Task |
|---|---|---|---|
| **7. SINGLE-LEG-STANCE RISE-TO-TOES EYES-CLOSED**<br>*SLS-RTT-EC*<br>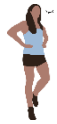<br>Single leg stance with rise-to-toe and eyes closed (5 seconds) on right and left leg. Hands on hips. | "We are going to add closing your eyes. As before, I will say 'go' and you will lift your LEFT leg, then I'll say 'rise' and you will rise to your toes, then when I say 'close' you will close your eyes. If needed can put your heel down or foot but try to stay as high on your toe as possible and keep your eyes closed. Then I will tell you to stand in neutral, you can rest, open your eyes, and we will repeat the other direction." | Video record: 0s<br>"Arm Raise & Lower": 2s<br>"Hands on hips": 4s<br>"Go": 6s<br>"Rise": 8s<br>"Eyes closed": 9s<br>"Neutral, eyes open": 12s<br>"Go": 14s<br>"Rise": 16s<br>"Eyes closed": 17s<br>"Neutral eyes open": 20s<br>Stop recording: 22s | |
| **8. FUNCTIONAL REACH TEST**<br>*FRT*<br>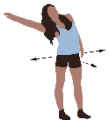<br>Forward and lateral (right and left), reaching to stability limit. Feet hip-width. | "Stand with your feet should width apart. This test is going to see how far you can reach without moving your feet. Follow my instructions - we will reach forward and then towards both sides. It is important to keep your heels on the ground." | Video record: 0s<br>"Arm Raise & Lower": 2s<br>"Reach forward": 5s<br>"Reach": 6s<br>"Reach": 8s<br>"Neutral": 9s<br>"Reach right": 11s<br>"Reach": 12s<br>"Reach": 14s<br>"Neutral": 15s<br>"Reach left": 17s<br>"Reach": 18s<br>"Reach": 20s<br>"Neutral": 21s<br>Stop recording: 23s | • Multi-Directional Reach Test (MDRT)<br>• Functional Reach Test<br>• BBS |
| **9. 5 TIMES SIT TO STAND**<br>*5TSTS*<br>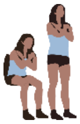<br>Five times sit to stand. Feet hip width. Arms crossed. | "Sit in the chair with your feet flat and your back against the back. Cross your arms over your chest. You are going to stand up and sit back down 5 times as fast as possible. Make sure you put weight into the chair each time and stand up all the way. You will finish sitting down." | Video record: 0s<br>"Arm Raise & Lower": 2s<br>"Go": 5s<br>"1, 2, 3, 4, 5": Count each repetition<br>Stop recording: 2s after returning to chair | • Five Times Sit to Stand Test<br>• Mini-BESTest<br>• SPPB<br>• BBS |

| Task | Instruction | Timing<br>(Recorded Cues) | Sample of Clinical Measures with Task |
|---|---|---|---|
| **10. Y-BALANCE**<br>*Y-Balance*<br>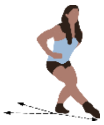<br>Reaching right and left foot in 3 directions: anterior, posteromedial, posterolateral, to stability limit. Stance foot flat. Hands on hips. | "I will walk you through this task while you perform it. Your goal in this exercise is to reach your foot as far as you can in different directions. For example, you will reach your right foot along the front line, then your left. Then we will reach each leg back along the diagonal to the side. And finally, back along the diagonal across your body. Keep your hands on your hips, keep your standing heel down, and keep your reaching foot just off the ground until you can't go further, then tap it on the ground." | Video record: 0s<br>"Arm Raise & Lower": 2s<br>"Hands on hips": 4s<br>"Go": 6s<br>"Reach right forward. Reach. Reach. Stand.<br>Reach left forward. Reach. Reach. Stand.<br>Reach right back diagonal. Reach. Reach. Stand.<br>Reach left back diagonal. Reach. Reach. Stand.<br>Reach right back across. Reach. Reach. Stand.<br>Reach left back across. Reach. Reach. Stand."<br>Stop recording: 2s after complete. | • Y-Balance Test<br>• Modified Star Excursion Balance Test (mSEBT) |
| **11. JUMP**<br>*Jump*<br>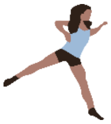<br>Jump as far as possible forward and laterally 45-degrees, facing forward, with double foot take-off to single-foot landing. Right (jump-r) and left (jump-l). | "Stand with two feet facing forward. Stay facing forward, jump in the forward diagonal direction and land on a single leg. If you are jumping to the right, land on your right leg. Try to stick your landing and stabilize as quickly as possible. You can use your arms however you'd like. Jump as far as you can while making sure you successfully land on one leg. We will do the right side and then stop and repeat on the left. Before we start, practice a jump on each side. | Video record: 0s<br>"Arm Raise & Lower": 2s<br>"Jump": 5s<br>"Hold, hold, hold, neutral": at landing<br>Stop recording: 2s after neutral | |
| **12. MARCH**<br>*March*<br>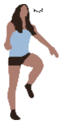<br>Marching in place with eyes closed. 50 steps. Arms straight with 45 degrees shoulder flexion. | "Stand at the center star and take a big step backwards. For this test we will have you extend your arms low to the front, close your eyes, and you will step or march in place for 50 steps. I will count for you and let you know when to stop so keep marching until I say stop." | Video record: 0s<br>"Arm Raise & Lower": 2s<br>"Arms low, eyes closed": 4s<br>"Go": 5s<br>"Neutral eyes open": After 50 steps - do not count out loud<br>Stop recording: 2s after neutral | • Fukuda Stepping Test<br>• Unterberger Step Test |

| Task | Instruction | Timing<br>(Recorded Cues) | Sample of Clinical Measures with Task |
|---|---|---|---|
| **13. GAIT**<br>*Gait*<br>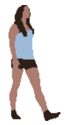<br>Walking forward at a natural pace, 3 meters. | "Start at the curtain. When I say 'go', walk forward along the line at a natural comfortable pace. Keep walking straight to the front camera, until I say stop. | Video record: 0s<br>"Arm Raise & Lower": 2s<br>"Go": 5s<br>Stop recording: once out of capture window | • SPPB<br>• Dynamic Gait Index (DGI)<br>• Functional Gait Assessment (FGA) |
| **14. GAIT-FAST**<br>*Gait-Fast*<br>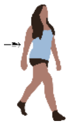<br>Walking with an increase in pace, from natural to 'fast' when cued (~1.5 meters) | "Start at the curtain. When I say go, begin walking at your normal speed, when I tell you 'fast', walk as fast as you can. When I say 'slow', walk very slowly. Keep walking straight until I say stop." | Video record: 0s<br>"Arm Raise & Lower": 2s<br>"Go": 5s<br>"Fast": 7s<br>Stop recording: once out of capture window | • Mini-BESTest<br>• DGI<br>• FGA |
| **15. GAIT-PIVOT**<br>*Gait-Pivot*<br>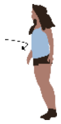<br>Natural pace gait with pivot turns to face opposite direction with feet together when cued to 'turn and stop'. | "Start at the curtain. When I say 'go', begin walking at your normal speed. When I tell you to 'turn and stop', turn as quickly as you can to face the opposite direction. After the turn, your feet should be together. When I say 'go', begin walking back until I say 'turn and stop' where you will again turn around with your feet close together. You can turn whichever direction you want, but make sure your feet are together after the turn." | Video record: 0s<br>"Arm Raise & Lower": 2s<br>"Go": 5s<br>"Turn and stop": at center star<br>"Go": after 2s<br>"Turn and stop": at final line<br>Stop recording: 2s after stop | • Mini-BESTest<br>• DGI<br>• FGA |
| **16. GAIT-TANDEM**<br>*Gait-Tandem*<br>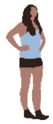<br>Tandem gait with dual task. Hands on hips. Gaze forward. Counting backwards by 3's. | "Start at the curtain. When I say 'go', walk with your feet aligned heel to toe in tandem like you're on a balance beam with your hands on your hips, looking forwards. Start at '94' and count backwards in 3's out loud. Try to continue stepping at the same pace, even if you slow your counting down." | Video record: 0s<br>"Arm Raise & Lower": 2s<br>"Go, 94": 5s<br>"Neutral": at end of line<br>Stop recording: 2 seconds after stop | • FGA (+ dual task) |

| Task | Instruction | Timing (Recorded Cues) | Sample of Clinical Measures with Task |
|---|---|---|---|
| **17. GAIT-BACKWARDS**<br>*Gait-Bwd*<br>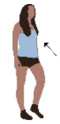<br>Walking backwards at a natural pace, 3 meters. | "Start at the front line. When I say 'go', walk backwards until I tell you to stop. For this task instead of a full arm raise at the beginning, you will raise your arm to the side just above shoulder height." | Video record: 0s<br>"Arm Raise & Lower": 2s<br>"Go": 5s<br>"Stop": past end of line<br>Stop recording: 2s after stop | • FGA |
| **18. GAIT EYES-CLOSED**<br>*Gait-EC*<br>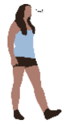<br>Walking with eyes-closed, 3 meters. | "Start at the curtain. When I say 'go' walk at your normal speed with your eyes closed." | Video record: 0s<br>"Arm Raise & Lower": 2s<br>"Go, eyes closed": 5s<br>"Stop": once out of capture window, before hitting tripods<br>Stop recording: 2s after stop | • FGA |

**Supplementary Table 2: StaBLE task goals and engineered features.**

| Task | Goal | Features |
|---|---|---|
| **1. STANDING-EC** | Stand still (explicit) | • Hold time, eyes open/closed (s)<br>• Sway (mean COM speed, final 5s), eyes open/closed (m/s)<br>$COM\ speed = \sqrt{v_{COM,AP}^2 + v_{COM,ML}^2}$<br>• Sway (C7 marker position standard deviation, final 5s), eyes open/closed (m/m)<br>$C7\ Sway = \frac{\sqrt{\sigma_{C7,AP}^2 + \sigma_{C7,ML}^2}}{h_{C7,t=0}}$ |
| **2. RISE-TO-TOES** | Rise as high as you can (explicit)<br>Rise as fast as you can (explicit)<br>Hold (explicit) | • Peak plantar flexion angle (mean right & left) (deg)<br>• Peak COM vertical speed during rise (m/s)<br>• Time COM is above 50% peak COM vertical position (s) |
| **3. SEMI-TANDEM** | Hold position (explicit)<br>As still as possible (explicit) | • Hold time, right/left (s)<br>• Sway (mean COM speed), right/left (m/s)<br>$COM\ speed = \sqrt{v_{COM,AP}^2 + v_{COM,ML}^2}$<br>• Sway (C7 marker position standard deviation), right/left (m/m)<br>$C7\ Sway = \frac{\sqrt{\sigma_{C7,AP}^2 + \sigma_{C7,ML}^2}}{h_{C7,t=0}}$ |
| **4. TANDEM** | Hold position (explicit)<br>As still as possible(explicit) | • Hold time, right/left (s)<br>• Sway (mean COM speed), right/left (m/s)<br>$COM\ speed = \sqrt{v_{COM,AP}^2 + v_{COM,ML}^2}$<br>• Sway (C7 marker position standard deviation), right/left (m/m)<br>$C7\ Sway = \frac{\sqrt{\sigma_{C7,AP}^2 + \sigma_{C7,ML}^2}}{h_{C7,t=0}}$ |

| Task | Goal | Features |
|---|---|---|
| **5. SINGLE-LEG-STANCE EYES-CLOSED** | Keep foot off the ground (explicit)<br><br>Hold as still as possible (explicit) | • Hold time (i.e., failure when lifted foot touches the ground or standing foot moves), right/left, eyes open/closed (s)<br><br>• Sway (mean COM speed), right/left, eyes open/closed (m/s)<br>$COM\ speed = \sqrt{v_{COM,AP}^2 + v_{COM,ML}^2}$<br>• Sway (C7 marker position standard deviation), right/left, eyes open/closed (m/m)<br>$C7\ Sway = \frac{\sqrt{\sigma_{C7,AP}^2 + \sigma_{C7,ML}^2}}{h_{C7,t=0}}$ |
| **6. SINGLE-LEG-STANCE RISE-TO-TOES** | Keep foot off the group (explicit)<br><br>Rise as high as possible (explicit)<br><br>Hold still (explicit) | • Hold time (i.e., failure when lifted foot touches the ground or standing foot heel lowers to the ground), right/left (s)<br><br>• Peak plantar flexion angle, right/left (deg)<br><br>• Sway (mean COM speed), right/left (m/s)<br>$COM\ speed = \sqrt{v_{COM,AP}^2 + v_{COM,ML}^2}$<br>• Sway (C7 marker position standard deviation), right/left, (m/m)<br>$C7\ Sway = \frac{\sqrt{\sigma_{C7,AP}^2 + \sigma_{C7,ML}^2}}{h_{C7,t=0}}$ |
| **7. SINGLE-LEG-STANCE RISE-TO-TOES EYES-CLOSED** | Keep foot of the group (explicit)<br><br>Rise as high as possible (explicit)<br><br>Hold still (explicit) | • Hold time (i.e., failure when lifted foot touches the ground or standing foot heel lowers to the ground), right/left (s)<br><br>• Peak plantar flexion angle, right/left (deg)<br><br>• Sway (mean COM speed), right/left (m/s)<br>$COM\ speed = \sqrt{v_{COM,AP}^2 + v_{COM,ML}^2}$<br>• Sway (C7 marker position standard deviation), right/left (m/m)<br>$C7\ Sway = \frac{\sqrt{\sigma_{C7,AP}^2 + \sigma_{C7,ML}^2}}{h_{C7,t=0}}$ |
| **8. FUNCTIONAL REACH TEST** | Reach as far as you can (explicit) | • Anterior reach distance, normalized by height (m/m)<br>$\frac{\max(x_{wrist} - x_{toe}), mean(R, L)}{height}$<br>• Right lateral reach distance, normalized by height (m/m)<br>$\frac{\max(z_{r,wrist} - z_{r,calc})}{height}$<br>• Left lateral reach distance, normalized by height (m/m)<br>$\frac{\max(z_{l,wrist} - z_{l,calc})}{height}$ |

| Task | Goal | Features |
|---|---|---|
| **9. 5 TIMES SIT-TO-STAND** | As fast as possible (explicit) | • Speed (m/s) $\frac{(\max(y_{COM}) - \min(y_{COM})) * 5}{total\ time}$ |
| **10. Y-BALANCE** | Reach foot as far as possible (explicit) | • Anterior reach distance, normalized by leg length, right/left (m/m)<br>• Posterolateral reach distance, normalized by leg length, left/right (m/m)<br>• Posteromedial cross reach distance, normalized by leg length, right/left (m/m) $\frac{\max(\sqrt{(x_{r,toe} - x_{l,toe})^2 + (z_{r,toe} - z_{l,toe})^2})}{\max(\lvert \overrightarrow{p}_{PSIS}^{calc} \rvert)}$ |
| **11. JUMP** | Jump as far as possible (explicit)<br><br>Hold your landing on a single leg (explicit) | • Jump distance, right/left (m/m) $\frac{\sqrt{(x_{calc,TD} - x_{calc,LO})^2 + (z_{calc,TD} - z_{calc,LO})^2}}{\text{height}}$ TD: touchdown, LO: liftoff<br>• Hold time before lifted foot touch down, right/left (s)<br>• Hold time before standing foot moves, right/left (s) |
| **12. MARCH** | Maintain original position and orientation (implicit) | • COM net translation (magnitude), global x-direction (m)<br>• COM net translation (magnitude), global z-direction (m)<br>• Pelvis net rotation (magnitude) (deg) |
| **13. GAIT** | Walk at natural pace (explicit) | • Mean COM speed (m/s) $mean(v_{x,COM})\rvert_{x_{COM}=0.5}^{x_{COM}=3.0}$ |
| **14. GAIT-FAST** | Increase gait speed at 'fast' cue (explicit) | • Change in COM speed (m/s) $mean(v_{x,COM})\rvert_{t_{start}}^{t_{start}+0.5} - mean(v_{x,COM})\rvert_{t_{end}}^{t_{end}+0.5}$ |
| **15. GAIT-PIVOT** | Turn as quickly as possible (explicit)<br><br>Keep feet together after turn (explicit) | • Mean pelvis rotation speed during turn (deg/s) $\lvert mean(\dot{q}_{pelvis\ rotation})\rvert \rvert_{\lvert q_{pelvis\ rotation}\rvert=20\ deg}^{\lvert q_{pelvis\ rotation}\rvert=160\ deg}$<br>• Ankle distance range after turn (m) $(max(ankle\ dist.) - min(ankle\ range))\rvert_{turn\ end\ (\lvert q_{pelvis\ rot.}\rvert=170\ deg)}^{gait\ start}$ $ankle\ dist. = \sqrt{(x_{r,calc} - x_{l,calc})^2 + (z_{r,calc} - z_{l,calc})^2}$ |

| Task | Goal | Features |
| --- | --- | --- |
| **16. GAIT-TANDEM** | Maintain pace (explicit)<br><br>Minimize sway (implicit) | • Mean COM forward speed (m/s) $mean(v_{x,COM})\|_{x_{COM}=0.5}^{x_{COM}=2.5}$<br><br>• Standard deviation of C7 position in mediolateral direction (m) $\sigma_{C7,ML}$ |
| **17. GAIT-BACKWARDS** | Walk at natural pace (implicit) | • Mean COM backward speed (m/s) $-mean(v_{x,COM})\|_{x_{COM}=2.5}^{x_{COM}=0.5}$ |
| **18. GAIT-EYES-CLOSED** | Maintain straight trajectory (implicit) | • Net mediolateral COM deviation (m) $\|((z_{COM})\|_{x_{COM}=3} - (z_{COM})\|_{x_{COM}=0})\|$ |

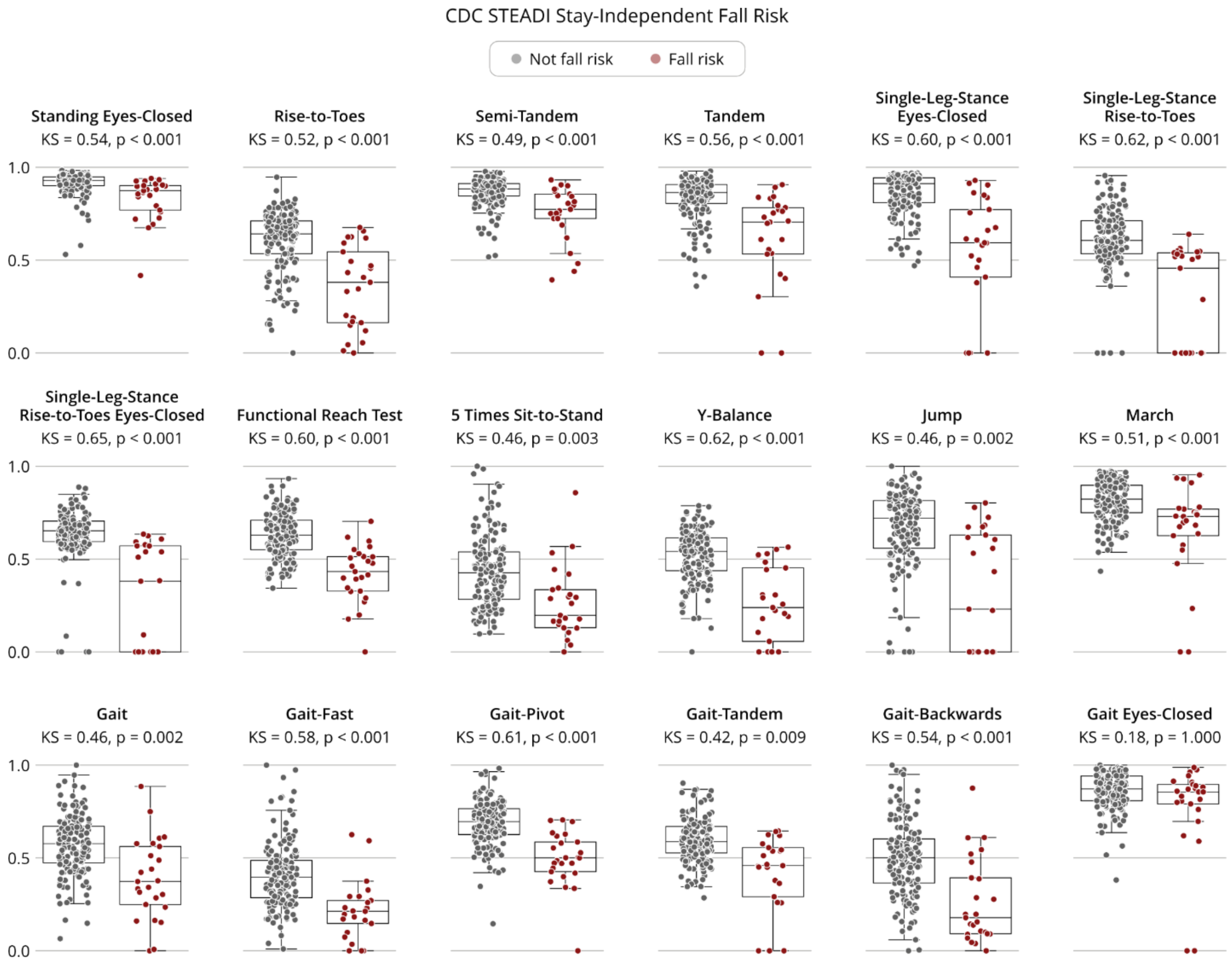


**Supplementary Fig. 1 | StaBLE task scores and fall risk.** Comparison of distributions of task scores for participants identified as fall-risk (CDC Stay-Independent 12-question tool score of 4 or greater, n = 25) and not at risk. Differences in underlying distributions are compared with the Kolmogorov–Smirnov test, applying the Bonferroni correction for multiple hypothesis testing.
